# Automated curation of CNMF-E-extracted ROI spatial footprints and calcium traces using open-source AutoML tools

**DOI:** 10.1101/2020.03.13.991216

**Authors:** LM Tran, AJ Mocle, AI Ramsaran, AD Jacob, PW Frankland, SA Josselyn

## Abstract

In vivo 1-photon calcium imaging is an increasingly prevalent method in behavioural neuroscience. Numerous analysis pipelines have been developed to improve the reliability and scalability of pre-processing and ROI extraction for these large calcium imaging datasets. Despite these advancements in pre-processing methods, manual curation of the extracted spatial footprints and calcium traces of neurons remains important for quality control. Here, we propose an additional semi-automated curation step for sorting spatial footprints and calcium traces from putative neurons extracted using the popular CNMF-E algorithm. We used the automated machine learning tools TPOT and AutoSklearn to generate classifiers to curate the extracted ROIs trained on a subset of human-labeled data. AutoSklearn produced the best performing classifier, achieving an F1 score > 92% on the ground truth test dataset. This automated approach is a useful strategy for filtering ROIs with relatively few labeled data points, and can be easily added to pre-existing pipelines currently using CNMF-E for ROI extraction.

## 1 Introduction

Advances in one-photon (1p) miniaturized fluorescence microscopy in terms of utility, cost, and ease-of-use have increased the accessibility and popularity of *in vivo* calcium imaging in freely behaving rodents (Cai et al., 2016; Ghosh et al., 2011; Hamel et al., 2015; Jacob et al., 2018). Consequently, researchers are able to track the activity of neuronal populations across days, weeks, or even months (Gonzalez et al., 2019; Rubin et al., 2015). Concurrent with the growing usage of 1p microendoscopy in neuroscience, there is an increasing demand for high-throughput software that accurately and efficiently processes the very large raw calcium imaging datasets now being produced. To address this challenge, a number of algorithms and analysis pipelines have been developed to automate the extraction of cells and calcium activity traces across time in a robust manner―a necessary step for downstream analyses (Pnevmatikakis, 2019).

Motion correction, source extraction, and cell registration (across multiple recording sessions) are important steps in pre-processing raw 1p calcium imaging data. Source extraction, the task of identifying the locations and activity of neurons in the imaged field of view (FOV), is arguably the most challenging of these steps is arguably the most challenging of these steps, as evidenced by the number of different algorithms published with the aim of improving this critical step. Nevertheless, two main methods of source extraction have been widely adopted in the field: principal component analysis/independent component analysis (PCA/ICA) (Mukamel et al., 2009) and the more recent extended constrained non-negative matrix factorization for microendoscopic data (CNMF-E) (Zhou et al., 2018). CNMF-E explicitly models background signals present in 1p microendoscopic recordings, and, therefore results in more accurate signal detection from neurons compared to PCA/ICA (Zhou et al., 2018).

Our lab has successfully applied CNMF-E to recordings from our open-source Compact Head-mounted Endoscope (CHEndoscope) in order to identify neuron locations (or spatial footprints) and extract their calcium activity traces from freely-behaving mice performing different behavioural tasks. CNMF-E has proven to be a reliable tool across multiple imaging sessions and experimental paradigms conducted in the lab with minimal parameter tuning in our hands (Jacob et al., 2018). However, like PCA/ICA, CNMF-E may still produce some false-positives in the output of detected cells (i.e., non-neuronal spatial footprints or calcium traces), which can be filtered out of the final dataset manually. We initially found success in filtering CNMF-E-extracted spatial footprints and traces by adding a manual curation step that involved visual inspection of each ROI and calcium trace (previously described in (Jacob et al., 2018)). While this type of manual curation can reduce the number of false-positives in CNMF-E’s output, visual inspection of potentially tens of thousands of extracted cells can be time-consuming, and this method is not free from human error. Here, we propose an automated machine learning (AutoML) approach built on top of the CNMF-E algorithm’s outputs to filter out potential false-positives. We implemented a semi-automated classification tool to limit the amount of manual curation required during pre-processing, without completely removing the ability to fine-tune the process with human-labeled datasets.

The main outputs of CNMF-E’s source extraction algorithm are: 1) the extracted calcium traces representing cellular activity, and, 2) the spatial footprint of putative neurons. As mentioned previously, manual curation of these outputs involves identifying both aberrant traces that do not have stable baseline fluorescence (Resendez et al., 2016), transient durations inconsistent with the expressed calcium indicator (e.g., GCAMP6f) (Badura et al., 2014), and/or spatial footprints that are not consistent with the shape and size of neurons in the brain region being recorded (Resendez et al., 2016). We trained and validated our classifiers on a dataset of 14 000 manually curated spatial footprints and traces output from CNMF-E. The final model chosen was then used to automate the curation of ROIs from other recording sessions. From the two AutoML libraries, we chose the best performing model to train on the full training set to evaluate on the test set. We find our model can accurately predict whether a cell would be included or excluded at a rate of 92%.

The potential time savings of manually curating thousands of cells makes this approach a method worth employing as part of a typical 1p calcium imaging pipeline. While our AutoML-based curation pipeline was primarily developed to be used with CHEndoscope data, our model takes the output of CNMF-E and as a result, allows this method to be readily applied to data acquired using other 1p miniature endoscopes.

## 2 Methods

### Dataset preparation and pre-processing

The dataset used for model training was acquired from multiple hippocampal CA1 recordings captured across different mice and recording sessions using methods described in Jacob et al. 2018. From these recordings, we used CNMF-E (Zhou et al., 2018) to extract spatial footprints and calcium traces of 14 000 ROIs. We then manually reviewed and labeled these ROIs as neuronal (included for further analysis) or artefact (excluded from analysis). The labels were generated by two human expert raters that inspected the calcium transients and spatial footprints based on previously reported criteria:

1. fast rise and slow decay of calcium transients with stable baseline fluorescence (Resendez et al., 2016).
2. calcium transient durations consistent with GCaMP6f (or appropriate GCaMP variant) (Badura, Sun, Giovannucci, Lynch, & Wang, 2014).
3. spatial footprints consistent with appropriate neuronal shape and size (Resendez et al., 2016).

Interrater agreement for the dataset was 87% across the two raters on a subset of the data (1073 putative ROIs extracted from CNMF-E) (Figure 1).

**Figure 1.**
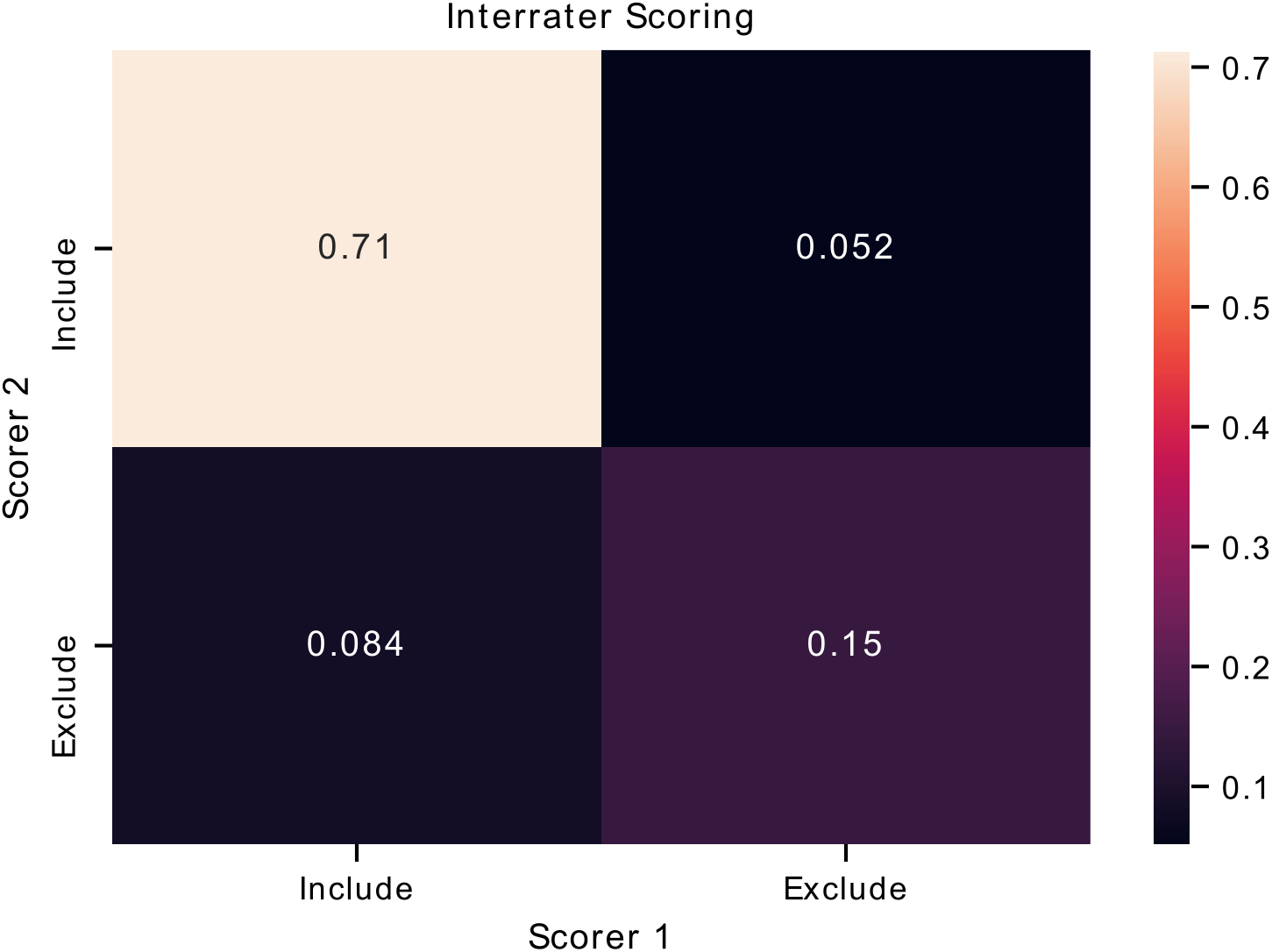
Interrater agreement of ROI labels. A confusion matrix comparing the manually reviewed labels (include of exclude) for putative ROIs extracted from CNMF-E determined by two different raters. Each cell of the matrix is annotated with the proportion of ROIs.

Spatial footprints consisted of the maximum projection of the identified cell from all frames in the video. We found that location of the footprint in the FOV was not important in our labelling criteria (compared to shape and size of footprint), we cropped the spatial footprints to remove empty space. Each spatial footprint was reduced to an 80×80 pixel image centered on the peak intensity of the footprint. Furthermore, recordings were of varying lengths, so all trace data was cropped at 500 frames (equivalent to 100s of recording at 5fps). The 2 dimensional footprints were reduced to a 1 dimensional vector (6400 pixels) and concatenated to the trace data.

We aggregated the labeled ROIs into a dataset split into training and test sets, which comprised 80% (~11 000 ROIs) and 20% (~3 000 ROIs) of the data, respectively.

### Model optimization and selection

We used two automated machine learning (AutoML) methods, TPOT (Olson et al., 2016; Olson & Moore, 2019) and AutoSklearn (Feurer et al., 2019) that are based on the popular Python machine learning toolbox, scikit-learn (Pedregosa et al. 2011) to select optimal classification models. While other AutoML tools exist that may outperform the ones we chose (Truong et al., 2019), TPOT and AutoSklearn are both free open-source, and easy to use, making them accessible for labs to incorporate into their existing analysis pipelines.

The key advantage of AutoML tools such as TPOT and AutoSklearn is that they do the extensive work of finding the best type(s) of data transformation and models to build a pipeline for classifying the input data, as well as the hyperparameters associated with these steps. TPOT is a tree-based optimization tool that builds and optimizes machine learning pipelines using genetic programming (Olson et al., 2016; Olson & Moore, 2019). TPOT generates pipelines of pre-processing steps and classification models in order to maximize classification performance while prioritizing simpler pipelines. AutoSklearn performs algorithm selection and hyperparameter tuning using Bayesian optimization, meta-learning and ensemble construction (Feurer et al., 2019) and as a result, the final classifier is an ensemble of many different model types and their associated hyperparameters. We primarily used default TPOT and AutoSklearn parameters, with a max evaluation time for a single pipeline of 10 minutes, and a total search time of 2 days.

During training, we used 10-fold cross-validation using stratified folds that preserved the relative proportions of “include” and “exclude” labels (i.e., during each run of training, 9 of 10 folds were used for training, and the 10th fold was used to test the performance of the model). This process was repeated for all 10 folds, resulting in an averaged performance metric for the data. We optimized the models to maximize the F1 score, the harmonic average of precision and recall, where high precision indicates a low false positive rate, and high recall indicates a low false negative rate. In our dataset, a true positive is an extracted ROI that both the trained model and a “ground truth” human scorer define as suitable to be included for further analysis (i.e., it satisfies the three selection criteria listed above). A true negative is an extracted ROI that is excluded for further analysis by both the model and our ground truth scoring.

## 3 Results

To determine the efficacy of an AutoML approach for classification of CNMF-E extracted ROIs, we tested the ability of TPOT and AutoSklearn to build classifiers that can label the pre-processed spatial footprints and calcium traces of putative ROIs. Both TPOT and AutoSklearn were trained on 11 000 labeled ROIs in the training set split into 10 folds for cross-validation, repeated 5 times. The best models obtained during training were used to determine the F1 score on the test set. Table 1 reports the performance of the best models obtained by TPOT and AutoSklearn across the training folds and on the test set.

**Table 1.** Mean F1 scores of AutoML methods on training cross-validation and final F1 scores on the test set.

|  | TPOT |  | AutoSklearn |  |
| --- | --- | --- | --- | --- |
|  | Training CV (10-fold, 5x) | Test | Training CV (10-fold, 5x) | Test |
| F1 Score | 0.906 | 0.904 | 0.917 | 0.922 |

Next we assessed the transferability of the best classifier pipelines identified by TPOT and AutoSklearn using fewer samples. We used the top performing classifier pipelines and hyperparameters chosen by TPOT and AutoSklearn and trained the initialized pipelines using datasets of increasing ROI number. The training set size ranged from 150 to 10 000 ROIs. Using a change point analysis algorithm (PELT, Killick et al. 2012), we determined that AutoSklean and TPOT classifiers approached a maximal F1 score with 719 and 1000 labeled ROIs, respectively (Figure 2). The pipelines found using our much larger labeled dataset may be easily incorporated into other pipelines with minimal computational effort to train and finetune on CNMF-E extracted ROIs from other 1p experiments, using fewer labeled ROIs.

**Figure 2.**
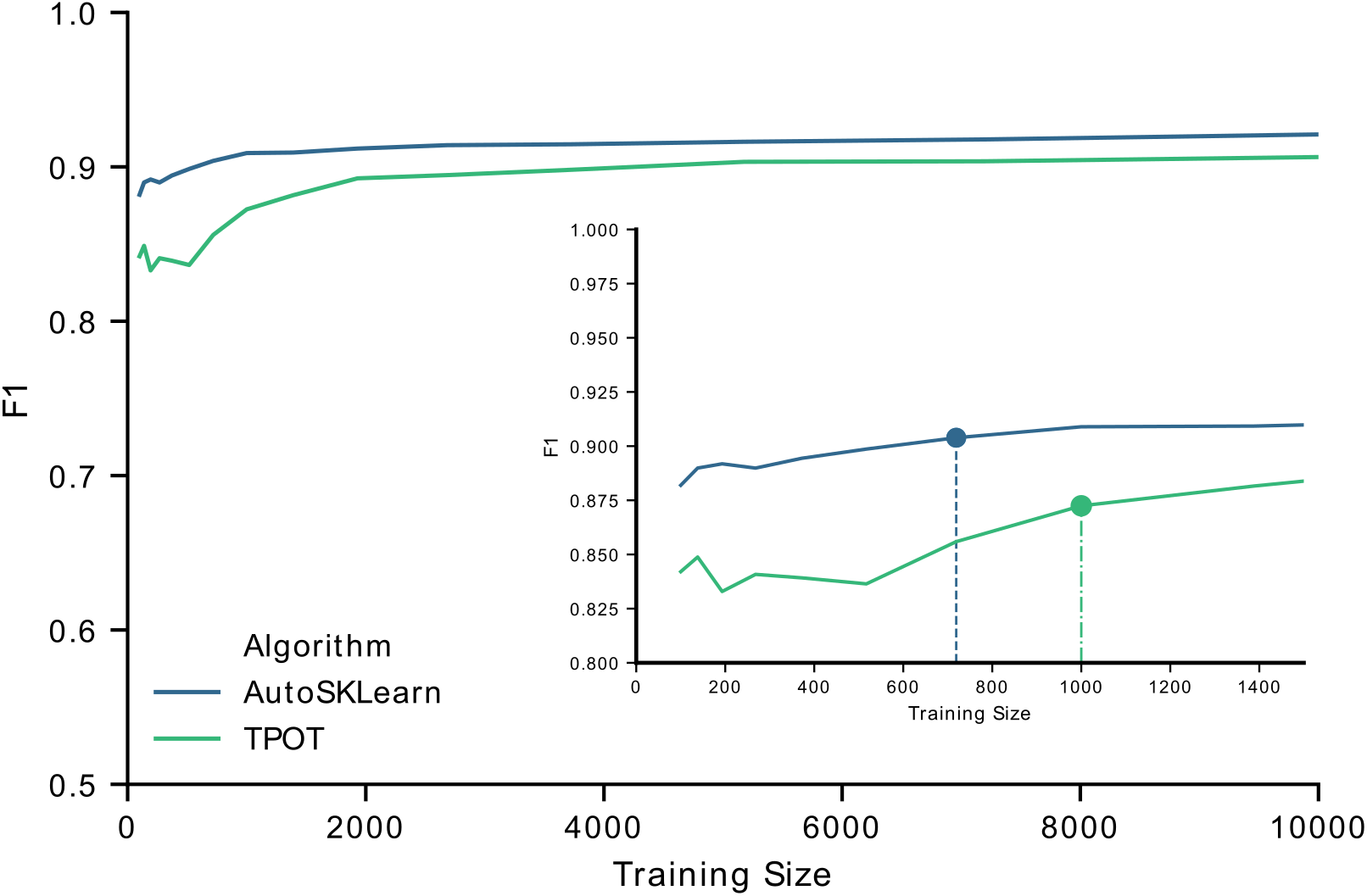
F1 score performance with increasing training size. Graph of the F1 test scores versus the number of training samples used to train the best models output by AutoSklearn (blue) and TPOT (green). (Inset) A graph of the same plot with a smaller range of training sizes and the change point is marked on each algorithm type.

To examine the classifier performance in terms of false positives and false negatives, we created confusion matrices to visualize the rate of true positives, true negatives, false positives and false negatives from the TPOT and AutoSklearn predictions compared to ground truth. We found that the classifier built with Autosklearn (0.922 F1, Table 1) performs better in terms of both reducing false positives and false negatives (Figure 3).

**Figure 3.**
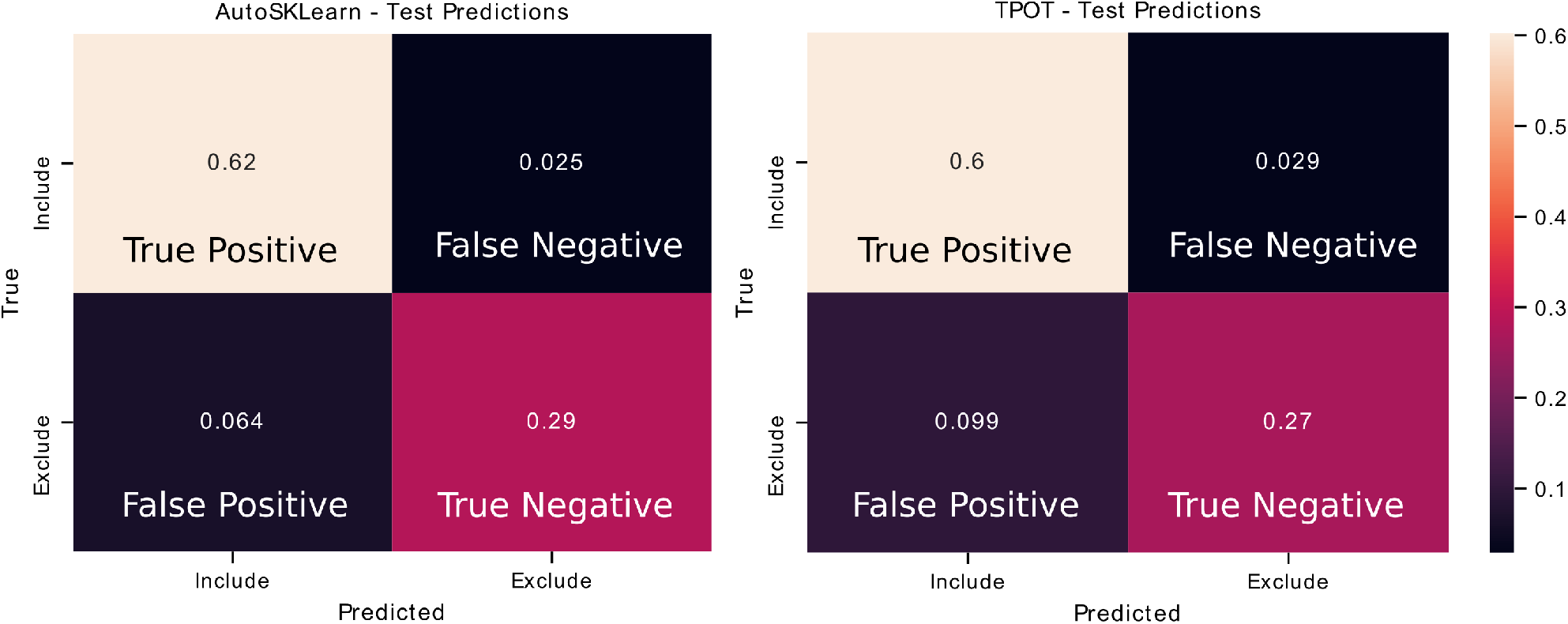
Confusion matrices of AutoML tools: TPOT and AutoSklearn. Each cell in the matrix is annotated with the proportion of ROIs labeled as Include or Exclude according to the predicted and true labels. Colors indicate the relative proportions of the labels where lower proportions are darker in color and higher proportions are lighter in color. The confusion matrices were made from predictions on the test set from the best models output by TPOT (left) and AutoSklearn (right).

To further assess the nature of the classification errors, we looked at the class confidences or probabilities of the test set predictions from the trained TPOT and AutoSklearn models (Figure 4). Class probabilities indicate the classifier’s certainty (using confidence score for TPOT and class probability for AutoSklearn) that a sample belongs to a particular class label. We tested whether mislabeled ROIs were also those which the classifiers expressed less confidence in classifying. We examined the size of the difference between certainty scores (true positives versus false positives, true negatives versus false negatives) in TPOT and AutoSklearn using Cohen’s d (Cohen, 2013; Sawilowsky, 2009) (Table 2). The AutoSklearn classifier―which outperformed the TPOT classifier based on F1 scores― showed large differences in certainty scores when labeling ROIs as positive (d=1.36) or negative (d=2.34). By contrast, the TPOT classifier was relatively less confident on both types of classification (positive d=0.63, negative d=1.68). In other words, the AutoSklearn classifier was more certain in applying labels to ROIs than was the TPOT classifier. This indicates that false negatives and false positives in the higher performing AutoSklearn classifier may arise from “edge-cases” ROIs in the dataset which the classifier was not as certain about the label. In contrast, the poorer performance of the TPOT classifier may simply be due to poor generalization on the test set.

**Table 2.** Cohen’s *d* of certainty scores between predicted labels that were correct or incorrect compared to ground truth in TPOT or AutoSklearn.

|  | Include (Positive) |  | Exclude (Negative) |  |
| --- | --- | --- | --- | --- |
|  | TPOT | AutoSklearn | TPOT | AutoSklearn |
| Cohen's $d$ | 0.63 | 1.36 | 1.68 | 2.34 |

**Figure 4.**
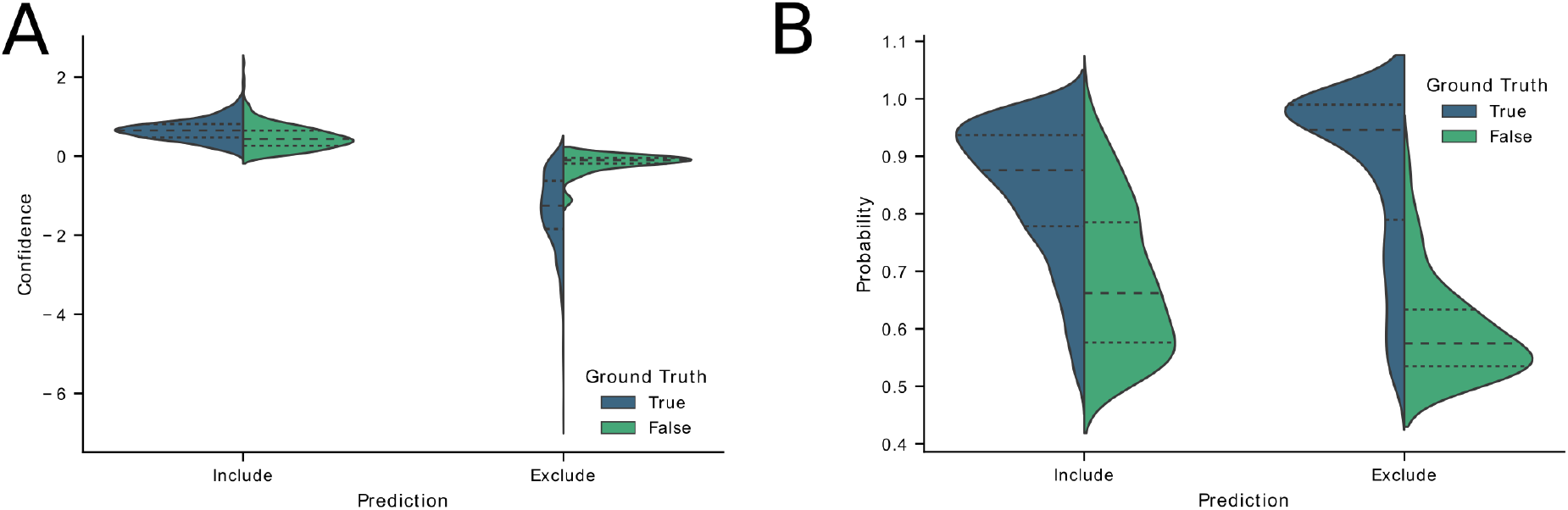
Classifier confidence (TPOT) and class probabilities (AutoSklearn) on predicted false positives and false negatives. Violin plots of the distribution of (A) TPOT classifier confidence or (B) AutoSklearn class probabilities on predicted ROI labels (Include or Exclude) in the test set. Each half of the violin plot is the distribution of values for labels that were correct (True, left/blue) or incorrect (False, right/green) based on the ground truth labels.

To investigate the nature of the false positives and false negatives from the best TPOT and AutoSklearn models, we looked at the underlying spatial footprints and calcium traces for mislabeled ROIs from both AutoML tools (Figure 5). Representative examples of excluded ROIs from the ground truth dataset show that some cells may be excluded (i.e., true negatives) because of poor trace data and/or poor spatial footprints, which possibly represent non-neuronal imaging artefacts and/or ROIs representing areas of background fluorescence. While some false positives from AutoSklearn shared similar features with true negative ROIs, others were more ambiguous. Upon inspection, these ROIs sometimes were high-quality spatial footprints with poor-quality calcium traces, or vice-versa, or were composed of spatial footprints and calcium traces of true neuronal-origin mixed with additional non-neuronal noise. These examples represent “edge-cases” which may be difficult to judge even by a human rater.

**Figure 5.**
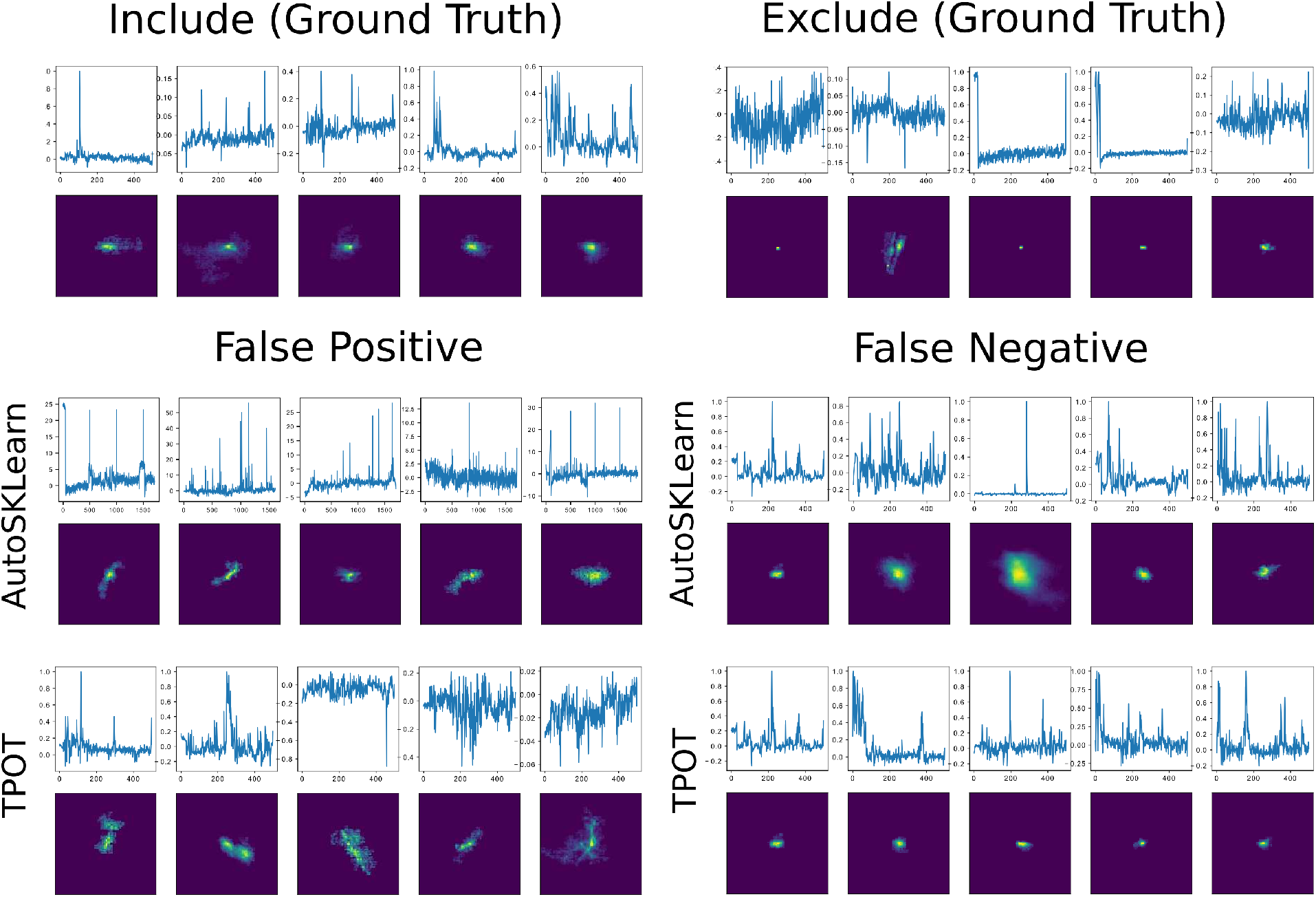
Representative false positives and negatives compared to ground truth ROIs. Example calcium traces (top) and spatial footprints (bottom) from ground truth ROIs labeled as Include (left) or Exclude (right). Example calcium traces (top) and spatial footprints (bottom) of false positive and false negative ROIs predicted from the AutoSklearn (middle row) or TPOT (bottom row) classifiers.

## 4 Discussion

Automated curation of ROIs provides a fast, accurate method for classifying neural data generated in 1p calcium imaging experiments. We show that AutoML tools such as the open source TPOT and AutoSklearn packages provide an easy way to build effective classifiers for ROIs extracted from the widely used CNMF-E algorithm. Spatial footprints and calcium traces from CNMF-E can be used to train these models with minimal data preprocessing. Furthermore, it may be possible to apply the top performing classifiers generated from this work to other experimental datasets taken from different 1p imaging setups, while requiring relatively few labeled samples. Other analyses pipelines such as MIN1PIPE (Lu et al. 2018) have been developed to improve source extraction by reducing false positive ROIs without increasing the rate of false negatives. However, given the more widespread adoption of CNMF-E, the approach described here prevents labs from having to adopt entirely new analysis pipelines. Our approach provides a balance between the need to manually review the output of CNMF-E ROIs to maximize the number of detected cells, while still allowing some further automation of the otherwise laborious curation process.

An AutoML approach to reviewing these traces may be useful for curating traces from labeled data of the extracted ROIs from CNMF-E and can be implemented on top of pre-existing analysis pipelines without much need to adapt the software. However, there are a number of limitations to this approach. Firstly, we emphasize the automated aspect of this machine learning classifier approach and little need for hand-tuning, but we recognize that the best models still make errors. Cases in which the best performing classifier generated by AutoSklearn failed to detect true positives or true negatives were further reviewed and were typically seen to be edge cases where it may be difficult for a human reviewer to make a judgment. Similarly, we found that a second expert scorer looking at the same data may not make the exact same judgments on such edge cases (having an interrater reliability score of 87%). While the AutoML classifiers were trained on the data that had relatively little preprocessing beyond cropping and downsampling, future work could address whether feature engineering over the spatial footprints and trace data could further improve accuracy and reduce training time for model selection and hyperparameter tuning. Better curation of a training dataset for the models may help reduce ambiguous cases that make it difficult for a classifier to make accurate predictions.

In conclusion, we present here a demonstration and benchmark of an AutoML approach for curation of CNMF-E extracted ROIs. The methods described here can provide a flexible, free open-source, and easy-to incorporate curation step for other researchers using CNMF-E for source extraction of their 1p datasets, while requiring few changes to their existing analysis pipelines.

## 6 Conflict of Interest

The authors declare that the research was conducted in the absence of any commercial or financial relationships that could be construed as a potential conflict of interest.

## 7 Author Contributions

LT, PF, SJ contributed to the study design. AJ designed and constructed the CHEndoscopes. AR and AM conducted surgeries, behavior experiments, CNMF-E processing, and manual ROI labelling. LT performed all analyses using automated machine learning pipelines. LT, AM, AR, AJ performed the statistical analyses and wrote the paper. All authors discusses and commented on the manuscript.

## 8 Funding

This work was supported by grants from the Canadian Institutes of Health Research (CIHR, grant numbers FDN-388455 to SJ, FDN143227 to PF), Natural Science and Engineering Council of Canada (NSERCs to SJ and PF), CIFAR catalyst award (SJ, PF) and an NIH (NIMH, 1 R01 MH119421-01) (SJ, PF). LT was supported by a SickKids Research Training Centre Restracomp Fellowship, Natural Science and Engineering Council of Canada Scholarship (NSERC, PGSD), AM by an NSERC CGSD, AR by an NSERC CGSD and NIH (NIMH,1 F31 MH120920-01), AJ by a Canadian Open Neuroscience Platform Student Scholar Award (in partnership with Brain Canada).

## 9 Acknowledgments

Compute resources provided by Compute Ontario and Compute Canada (www.computecanada.ca) were used to perform this research.

## Data Availability Statement

The datasets and code generated for this study can be found in the cnmfe-reviewer GitHub repository [https://github.com/jf-lab/cnmfe-reviewer].

## Notes

https://github.com/jf-lab/cnmfereview

